# Metastatic signaling of hypoxia-related genes across TCGA Pan-Cancer types

**DOI:** 10.1101/2020.02.01.930479

**Authors:** Andrés López-Cortés, Patricia Guevara-Ramírez, Santiago Guerrero, Esteban Ortiz-Prado, Jennyfer M. García-Cárdenas, Ana Karina Zambrano, Isaac Armendáriz-Castillo, Andy Pérez-Villa, Verónica Yumiceba, Nelson Varela, Daniel Córdova-Bastidas, Paola E. Leone, César Paz-y-Miño

## Abstract

Many primary-tumor subregions have low levels of molecular oxygen, termed hypoxia. Hypoxic tumors are at elevated risk for local failure and distant metastasis. Metastatic disease is the leading cause of cancer-related deaths and involves critical interactions between tumor cells and the microenvironment. Here we focused on elucidating the molecular hallmarks of tumor hypoxia that remains poorly defined. To fill this gap, we analyzed the genomic alterations and hypoxia score of 233 hypoxia-related genes of 6,343 individuals across 17 TCGA Pan-Cancer types. In addition, we analyzed a protein-protein interactome (PPi) network and the shortest paths from hypoxic proteins to metastasis. As results, mRNA high alteration was prevalent in all cancer types. Genomic alterations and hypoxia score presented a highest frequency in tumor stage 4 and positive metastasis status in all cancer types. The most significant signaling pathways were HIF-1, ErbB, PI3K-Akt, FoxO, mTOR, Ras and VEGF. The PPi network revealed a strong association among hypoxic proteins, cancer driver proteins and metastasis driver proteins. The analysis of shortest paths revealed 99 ways to spread metastasis signaling from hypoxic proteins. Additionally, we proposed 62 hypoxic genes strongly associated with metastasis and 27 of them with high amount of genomic alterations. Overall, tumor hypoxia may drive aggressive molecular features across cancer types. Hence, we identified potential biomarkers and therapeutic targets regulated by hypoxia that could be incorporated into strategies aimed at improving novel drug development and treating metastasis.

## INTRODUCTION

The Nobel Prize in Physiology or Medicine for 2019 was awarded for the discoveries of how cells sense and adapt to oxygen availability. Human cells undergo fundamental shifts in gene expression when there are changes in the oxygen (O_2_) levels around them ^1^. These changes in gene expression alter tissue remodeling, cell metabolism, and even organismal responses such as increases in heart rate and ventilation. In 1990’s, the Hypoxia Inducible Factor (*HIF*) was identified, purified and cloned. *HIF* is a transcription factor that regulates these oxygen-dependent responses and consisted of two components: *HIF-1* and *ARNT* ^2–4^. In 1995, the von Hippel-Lindau (*VHL*) tumor suppressor gene was studied and the first full-length clone of the gene was isolated ^5–8^. Consequently, it was demonstrated in 1999 that *VHL* regulates *HIF-1* post-transcriptional and oxygen-sensitivity degradation ^9,10^.

Molecular O_2_ is a key nutrient required for aerobic metabolism to maintain intracellular bioenergetics and as a substrate in numerous organic and inorganic reactions. Hypoxia occurs in a variety of physiological as well as pathological conditions ^11^. Half of all solid tumors are characterized by dynamic gradients of O_2_ distribution and consumption, that is, the tumor environment presents hypoxia subregions due to changes in tumor metabolism that increases O_2_ demand or irregular tumor vasculature that decreases O_2_ supply ^12–18^. Tumor adaptation to this imbalance between O_2_ demand and supply is associated with increased genomic instability and poor clinical prognosis ^15,19^, development of tumor stem cell-protective niches ^20,21^, resistance to radio- and chemotherapy ^22,23^, and increased proclivity for distant metastasis ^24–26^.

Metastasis is thought to be the ultimate manifestation of a neoplastic cell’s evolution towards becoming autonomous from the host, and remains the main cause of cancer-related death ^27,28^. The persistence and lethal relapse of disseminated cancer is driven by stem-like cells that have the ability to regenerate tumors in distant sites ^29–32^. In fact, cure of most cancers is probable whenever diagnosis occurs before cells have spread beyond the tissue of origin; otherwise, cancer is often referred to as incurable ^33–35^. According to Welch and Hurst ^28^, the four hallmarks of metastasis are motility and invasion, modulate microenvironment, plasticity and colonization. Regarding mechanisms of HIF-mediated metastasis, hypoxia and activation of HIF signaling influence multiple steps within the metastatic cascade, including invasion and migration, intravasation and extravasation, and establishment of the premetastatic niche, as well as survival and growth at the distant site ^24,36,37^.

Although hypoxia is an adverse and targetable prognostic feature in multiple cancer types ^38,39^, both the protein-protein interactome (PPi) network and the shortest paths from hypoxia-related genes (HRG) to metastasis had not been described. To fill this gap, we evaluated 233 HRG in 6,343 tumors representing 17 different cancer types. We analyzed genomic alterations and Buffa hypoxia scores per tumor stage, metastatic status and ethnicity across multiple tumor types. We analyzed functional enrichment analysis of hypoxic genes. Lastly, we detailed all information about clinical trials.

## RESULTS

### OncoPrint of genomic alterations

We analyzed genomic alterations of 6,343 individuals with 17 different cancer types taken from the Pan-Cancer Atlas (PCA) project from The Cancer Genome Atlas (TCGA) ^40–49^. Figure 1A shows the OncoPrint of genomic alterations (mRNA high, mRNA low, copy number variant (CNV) deep deletion, CNV amplification, fusion gene, inframe mutation, truncating mutation and missense mutation) of the 233 hypoxia-related genes using cBioPortal ^50,51^. The TCGA Pan-Cancer types were bladder urothelial carcinoma (BLCA) with 369 individuals, breast invasive carcinoma (BRCA) with 991 individuals, cervical squamous cell carcinoma and endocervical carcinoma (CESC) with 217 individuals, colorectal adenocarcinoma (CRC) with 521 individuals, esophageal carcinoma (ESCA) with 163 individuals, head and neck squamous cell carcinoma (HNSC) with 431 individuals, kidney renal clear cell carcinoma (KIRC) with 352 individuals, liver hepatocellular carcinoma (LIHC) with 345 individuals, lung adenocarcinoma (LUAD) with 501 individuals, lung squamous cell carcinoma (LUSC) with 464 individuals, mesothelioma (MESO) with 80 individuals, pancreatic adenocarcinoma (PAAD) with 166 individuals, prostate adenocarcinoma (PRAD) with 481 individuals, skin cutaneous melanoma (SKCM) with 258 individuals, stomach adenocarcinoma (STAD) with 399 individuals, testicular germ cell tumors (TGCT) with 127 individuals and thyroid carcinoma (THCA) with 478 individuals.

**Figure 1.**
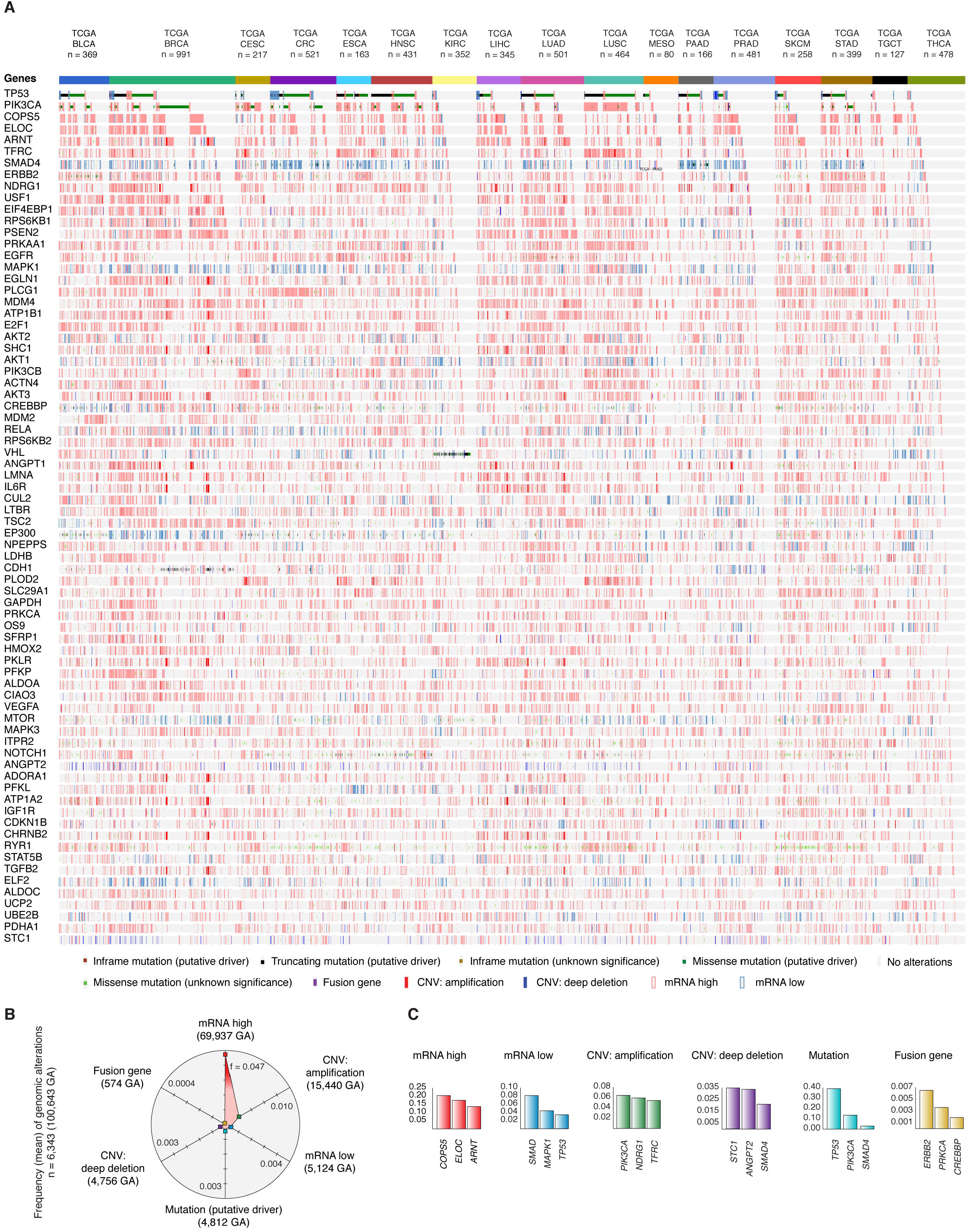
Panoramic landscape of genomic alterations across TCGA Pan-Cancer types. A) OncoPrint of genomic alterations (mRNA high, mRNA low, CNV amplification, CNV deep deletion, putative driver mutation and fusion genes) of the most altered hypoxia-related genes. B) Most frequency genomic alterations across all cancer types. C) Most altered genes per genomic alteration type.

We found 100,643 genomic alterations in 6,343 individuals. The genomic alteration with the highest frequency mean was mRNA high (0.047), followed by CNV amplification (0.010), mRNA low (0.004), putative driver mutation (0.003), CNV deep deletion (0.003) and fusion gene (0.0004) (Figure 1B and Table S1).

Consequently, genes with the highest frequency mean per genomic alteration were *COPS5* (0.204), *ELOC* (0.175), *ARNT* (0.137), *PIK3CA* (0.135) and *TFRC* (0.117) with mRNA high; *SMAD4* (0.081), *MAPK1* (0.044), *TP53* (0.034), *ELF2* (0.031) and *AKT1* (0.031) with mRNA low; *PIK3CA* (0.064), *NDRG1* (0.059), *TFRC* (0.054), *ANGPT1* (0.053) and *ARNT* (0.049) with CNV amplification; *STC1* (0.035), *ANGPT2* (0.034), *SMAD4* (0.021), *TEK* (0.016) and *VEGFC* (0.015) with CNV deep deletion; *TP53* (0.391), *PIK3CA* (0.134), *SMAD4* (0.029), *VHL* (0.026) and *CDH1* (0.023) with putative driver mutations; *ERBB2* (0.006), *PRKCA* (0.004), *TP53* (0.002), *CREBBP* (0.002) and *STAT3* (0.002) with fusion genes; finally, *TP53* (0.476), *PIK3CA* (0.341), *COPS5* (0.246), *ELOC* (0.213) and *ARNT* (0.198) with all genomic alterations (Figure 1C and Table S1).

### Genomic alterations per TCGA Pan-Cancer type

Figure 2A shows the frequency mean of genomic alterations per TCGA Pan-Cancer type (0.067). The top ten TCGA Pan-Cancer types with the highest frequency mean of genomic alterations were ESCA (0.089), BRCA (0.080), LUSC (0.080), BLCA (0.077), LUAD (0.076), STAD (0.072), LIHC (0.071), CESC (0.069), HNSC (0.067) and CRC (0.066) (Tables S2-S18).

**Figure 2.**
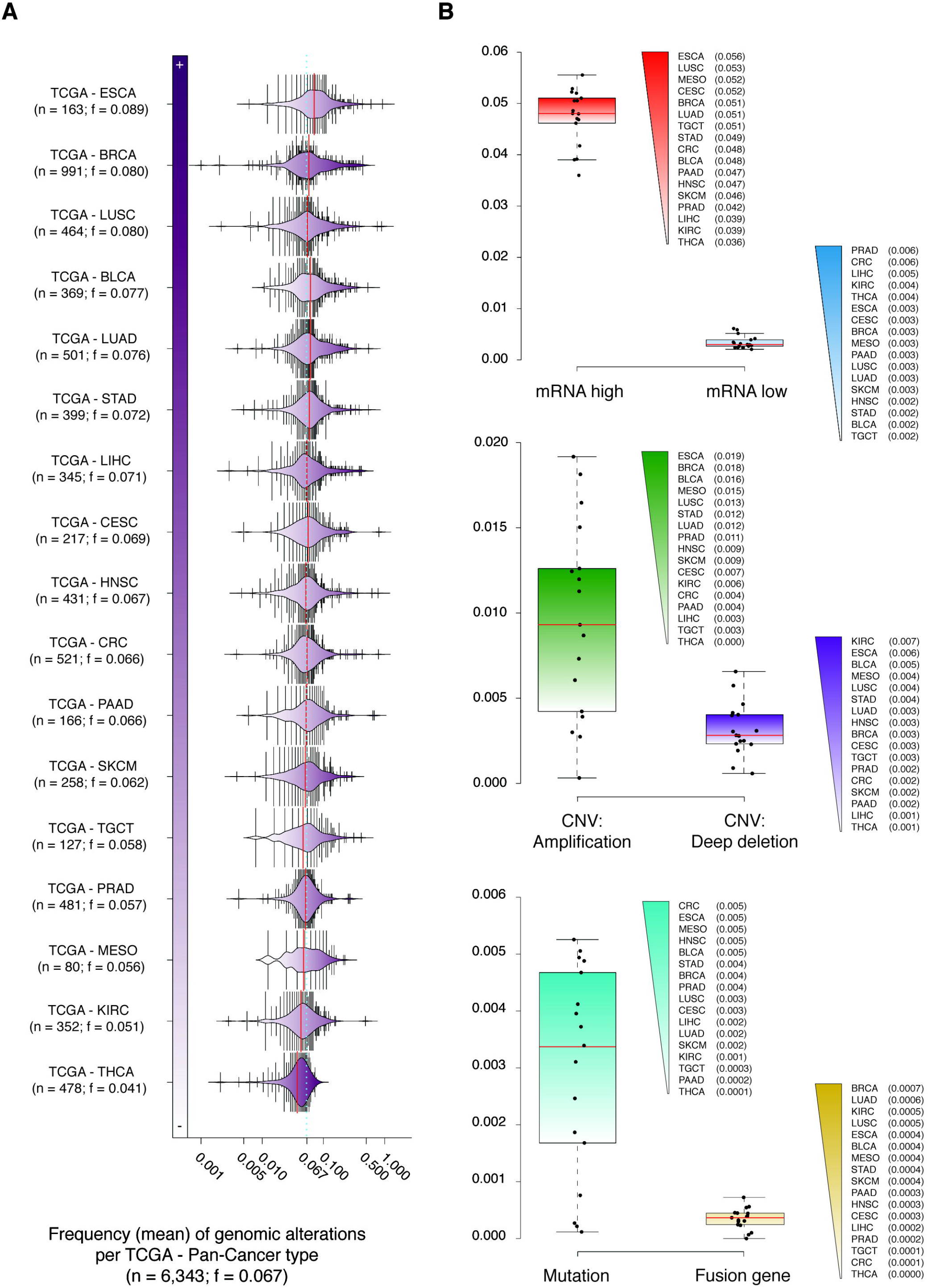
Genomic alterations per TCGA Pan-Cancer type. A) Frequency mean of genomic alterations per cancer type. B) Most altered cancer types per genomic alteration.

TCGA Pan-Cancer types with the highest frequency mean per genomic alteration were ESCA (0.056), LUSC (0.053) and MESO (0.052) with mRNA high; PRAD (0.006), CRC (0.006) and LIHC (0.005) with mRNA low; ESCA (0.019), BRCA (0.018) and BLCA (0.016) with CNV amplification; KIRC (0.007), ESCA (0.006) and BLCA (0.005) with CNV deep deletion; CRC (0.005), ESCA (0.005) and MESO (0.005) with putative driver mutations; and BRCA (0.0007), LUAD (0.0006) and KIRC (0.0005) with fusion genes (Figure 2B).

### Tumor stage and metastasis status

Figure 3A shows the frequency mean of genomic alterations of 17 TCGA Pan-Cancer types per tumor stage. The frequency mean of T1 stage was 0.060 (n = 1,396), of T2 stage was 0.072 (n = 2,306), of T3 stage was 0.068 (n = 2,029), and of T4 was 0.072 (n = 612) (Table S19-S22). The Mann-Whitney *U* test showed a significant difference of genomic alterations between T1 and T4 stages (P < 0.001).

**Figure 3.**
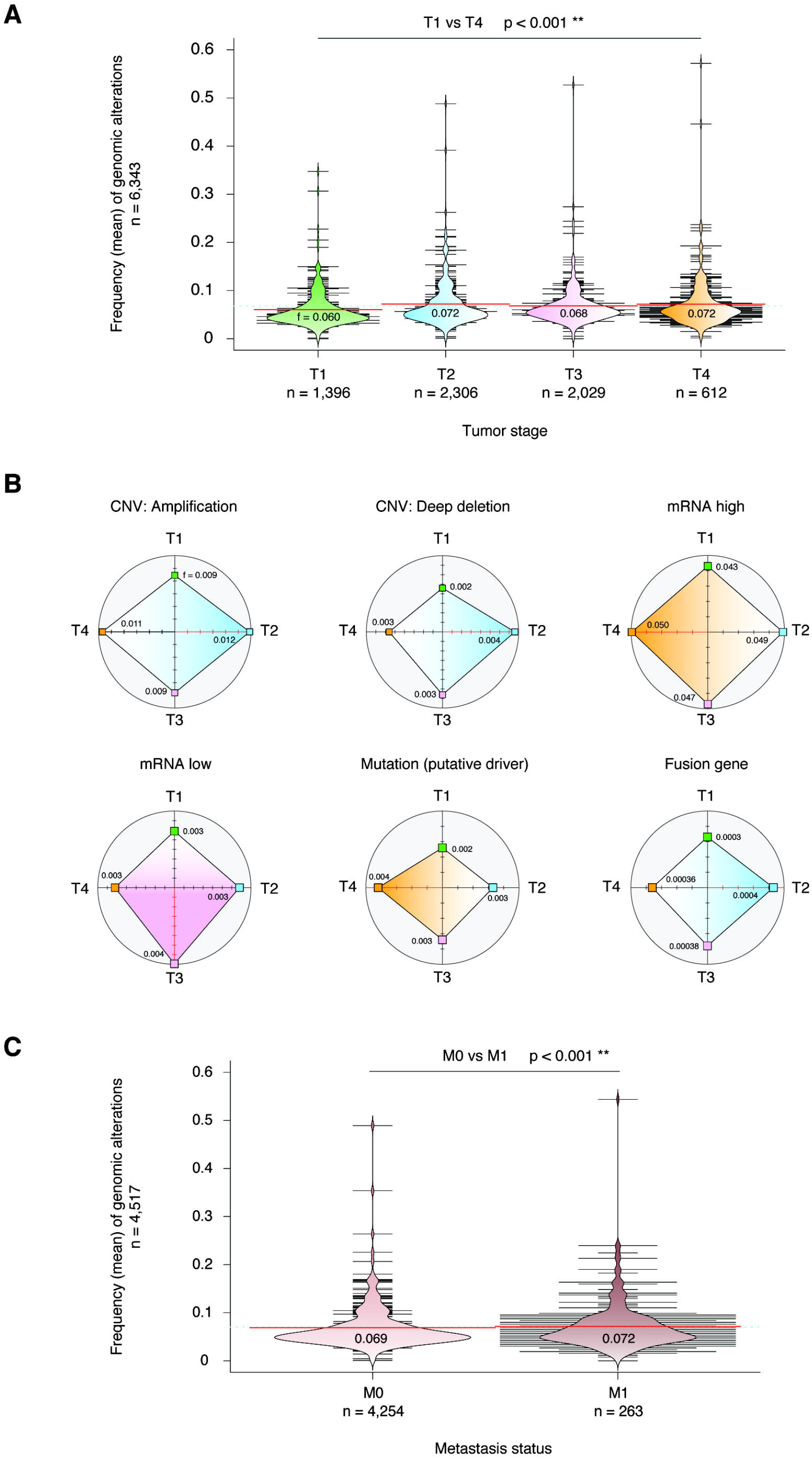
Tumor stages and metastasis status. A) Frequency mean of all genomic alterations per tumor stage across 17 Pan-Cancer types. B) Frequency mean of each genomic alteration per tumor stage. C) Frequency mean of all genomic alterations per metastasis status.

Figure 3B shows the association between genomic alterations and the tumor stage with the highest frequency mean. T2 presented the highest frequency mean of CNV amplifications (0.012), CNV deep deletions (0.004) and fusion genes (0.0004); T3 presented the highest frequency mean of mRNA low; and T4 presented the highest frequency mean of mRNA high (0.050) and putative driver mutations (0.004).

On the other hand, figure 3C shows the frequency mean of genomic alterations per metastasis status. The frequency mean of M0 was 0.069 (n = 4,254) and of M1 was 0.072 (n = 263) (Table S23). The Mann-Whitney *U* test showed a significant difference of genomic alterations between M0 and M1 (P < 0.001).

### Hypoxia score

We first quantified tumor hypoxia in 5,249 tumors from 13 tumor types in the PCA and TCGA by using Buffa signature ^52^. Figure 4A shows the mean of Buffa hypoxia score (HS) per cancer type. The most hypoxic TCGA Pan-Cancer types were HNSC (29.6), LUSC (26.8), CESC (21), CRC (17.3), BLCA (14.5), SKCM (5), KIRC (2.2), LUAD (−0.5), PAAD (−8.5) and BRCA (−9.6). The overall mean of HS in the 13 TCGA Pan-Cancer types was 0.85.

**Figure 4.**
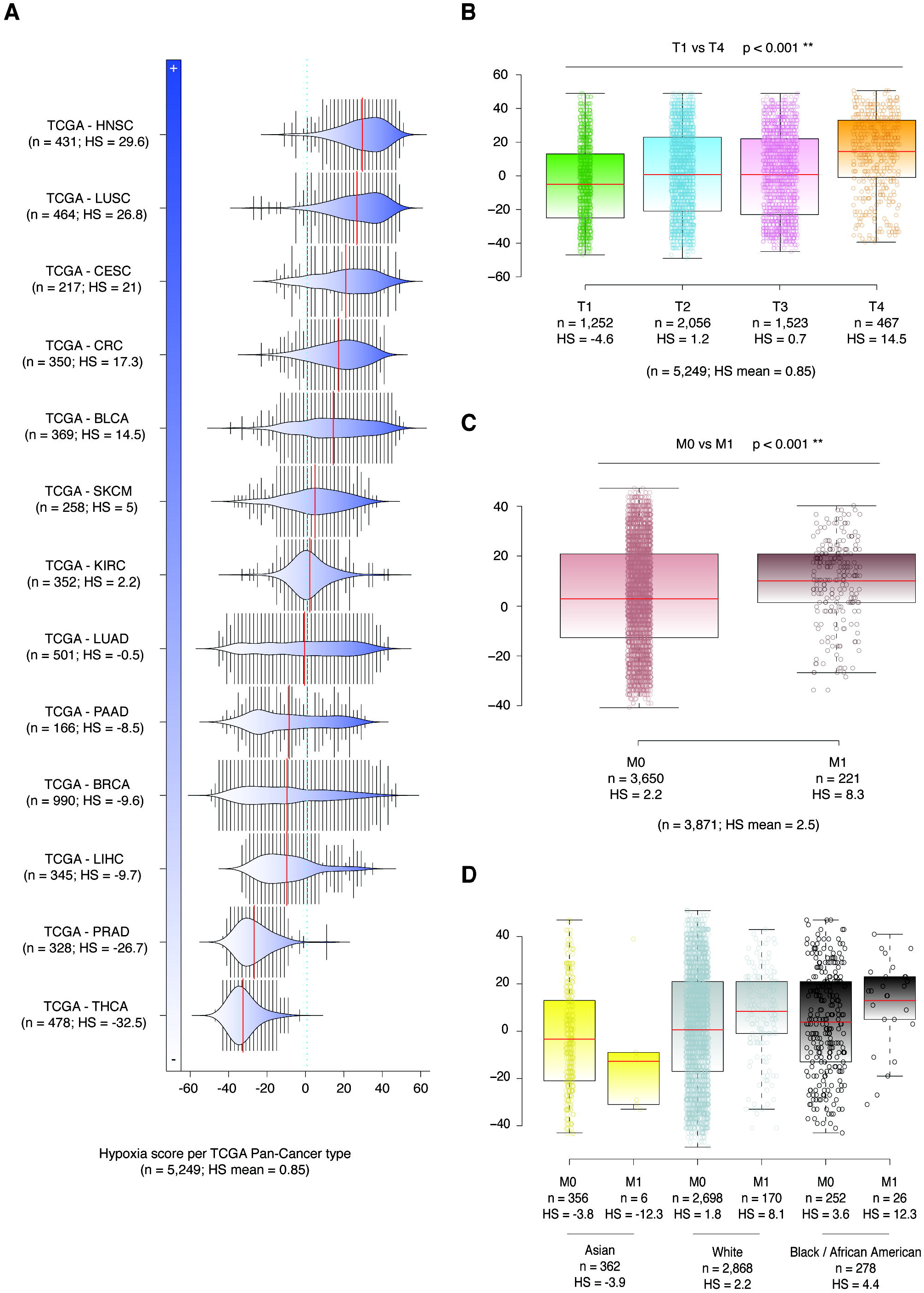
Hypoxia score. A) Hypoxia score mean across 13 Pan-Cancer types. B) Hypoxia score mean per tumor stage. C) Hypoxia score mean per metastasis status. D) Hypoxia score mean per metastasis status related to ethnicity.

Figure 4B shows box plots related to the Buffa HS mean per tumor stage (T1-T4) in 5,249 individuals with 13 different TCGA Pan-Cancer types. The HS mean in T1 was −4.6, in T2 was 1.2, in T3 was 0.7, and in T4 was 14.5. Consequently, we obtained a significant difference of hypoxia scores between T1 and T4 using the Mann-Whitney *U* test (P < 0.001).

Figure 4C shows box plots of Buffa HS in 3,871 individuals with metastasis status (M0 and M1). The HS mean in individuals without metastasis (M0) was 2.2, while in individuals with metastasis (M1) was 8.3. Consequently, we obtained a significant difference between the hypoxia scores of M0 and M1 (Mann-Whitney *U* test, P < 0.001).

Finally, figure 4D shows box plots of Buffa HS in 3,508 individuals with both ethnicity information and metastasis status. The HS mean in black / African Americans was 4.4, in white individuals was 2.2, and in Asians was −3.9. Regarding metastasis status per ethnic group, black/ African Americans had a HS mean of 3.6 in M0 and an increased 12.3 in M1, white individuals had 1.8 in M0 and an increased 8.1 in M1, and Asians had −3.8 in M0 and a decreased −12.3 in M1.

### Protein expression analysis

The Human Protein Atlas (HPA) presented a map of the human tissue proteome based on tissue microarray-based immunohistochemistry. HPA has analyzed 175 of 233 hypoxia-related proteins across 15 cancer types, classifying the protein expression in high, medium, low and non-detected ^53–55^. As results, AKT3, CAMK2A, PDHA1, PDHB, POLR2A, SMAD4 and SOD3 were proteins with high and medium expression in normal tissues, and non-detected expression in a consensus of 15 cancer types, acting as tumor suppression proteins. Meanwhile, ADAM8, BBC3, CCL2, CDH1, CLDN3, EDN1, EIF4EBP1, RELA and TNC were proteins with high and medium expression in a consensus of 15 cancer types, and non-detected expression in normal tissues, acting as oncogenes (Figure 5 and Table S24).

**Figure 5.**
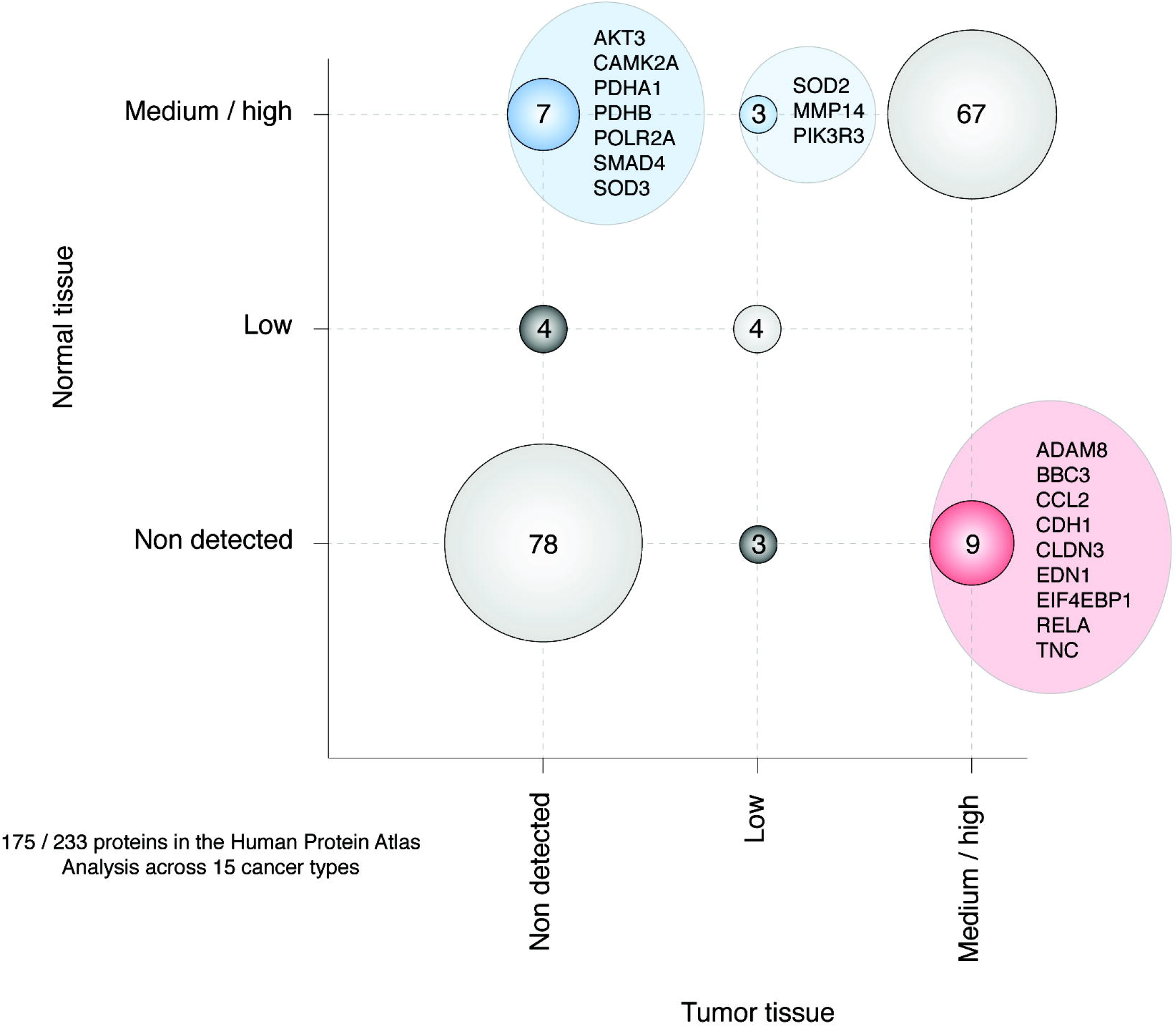
Molecular signatures of proteomes in the human tissue based on tissue microarray-based immunohistochemistry according to the Human Protein Atlas.

### Hallmarks of cancer

Figure 6 shows a circos plot of 23 (10%) HRG that are hallmarks of cancer according to Hanahan and Weinberg ^56^. The top 10 genes with the highest number of interactions with the hallmarks of cancer were *TP53*, *AKT1*, *EGFR*, *EPAS1*, *HIF1A*, *MTOR*, *NOTCH1*, *VHL*, *CREBBP* and *ERBB2*. Suppression of growth was promoted by *TP53*, *AKT1*, *EPAS1*, *VHL*, *NDRG1*, *SMAD3*, *PIK3R1*, and *EP300*. Escaping immune response to cancer was promoted by *EGFR* and *HIF1A*. Cell replicative immortality was promoted by *TP53* and *NOTCH1*. Tumor promoting inflammation was promoted by *EPAS1*. Angiogenesis was promoted by *AKT1*, *ARNT*, *EGFR*, *EPAS1*, *HIF1A*, *MTOR*, *NOTCH1*, *PIK3CA* and *PLCG1*. Escaping programmed cell death was promoted by *TP53*, *AKT1*, *EGFR*, *EPAS1*, *HIF1A*, *MTOR*, *NOTCH1*, *CREBBP*, *ERBB2*, *PIK3CA*, *PLCG1*, *SMAD3*, *MAPK1*, *ATP1A1* and *PIK3CB*. Change of cellular energetics was promoted by *TP53*, *AKT1*, *EGFR*, *HIF1A*, *MTOR*, *NOTCH1*, *VHL*, *ERBB2*, *ARNT*, *MAPK1* and *CDH1*. Proliferative signaling was promoted by *AKT1*, *EGFR*, *EPAS1*, *MTOR*, *NOTCH1*, *CREBBP*, *ERBB2*, *NDRG1*, *PIK3CA*, *PLCG1*, *SMAD3* and *CXCR4*. Lastly, invasion and metastasis was promoted by *TP53*, *AKT1*, *EGFR*, *EPAS1*, *HIF1A*, *MTOR*, *VHL*, *CREBBP*, *ERBB2*, *NDRG1*, *PIK3CA*, *PLCG1*, *SMAD3*, *ARNT*, *MAPK1*, *PIK3R1*, *ATP1A1*, *CDH1* and *CXCR4*.

**Figure 6.**
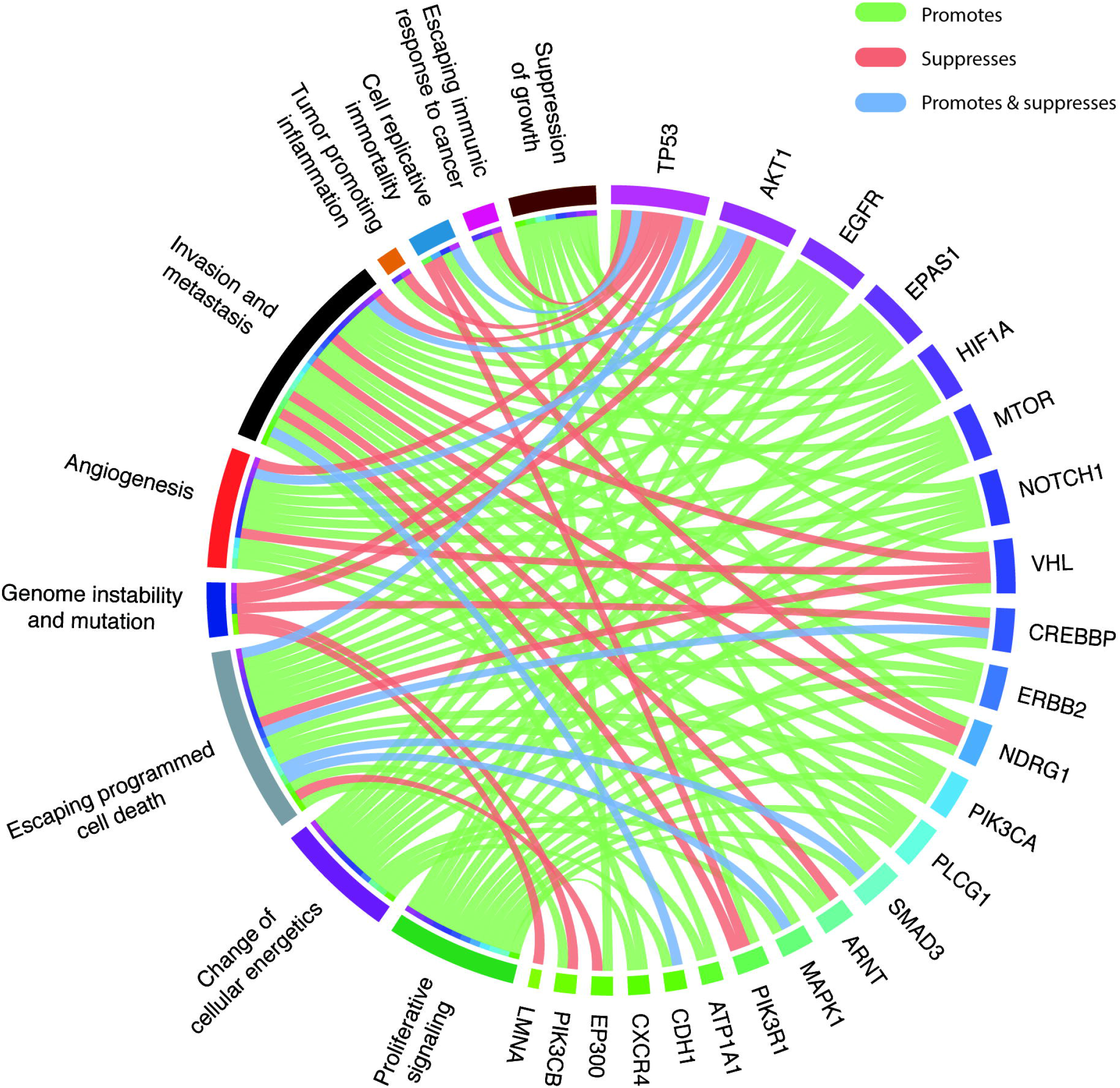
Hypoxia-related genes with hallmark of cancer signatures.

### Protein-protein interactome network

The PPi network was performed to better understand the behavior of hypoxia-related proteins in cancer and metastasis using the String Database and Cytoscape (Figure 7) ^57,58^. With the indicated cutoff of 0.9, the final interactome network had 285 nodes conformed by 163 (57%) hypoxia-related proteins, 98 (34%) cancer driver proteins, and 24 (8%) hypoxia-related proteins and cancer driver proteins.

**Figure 7.**
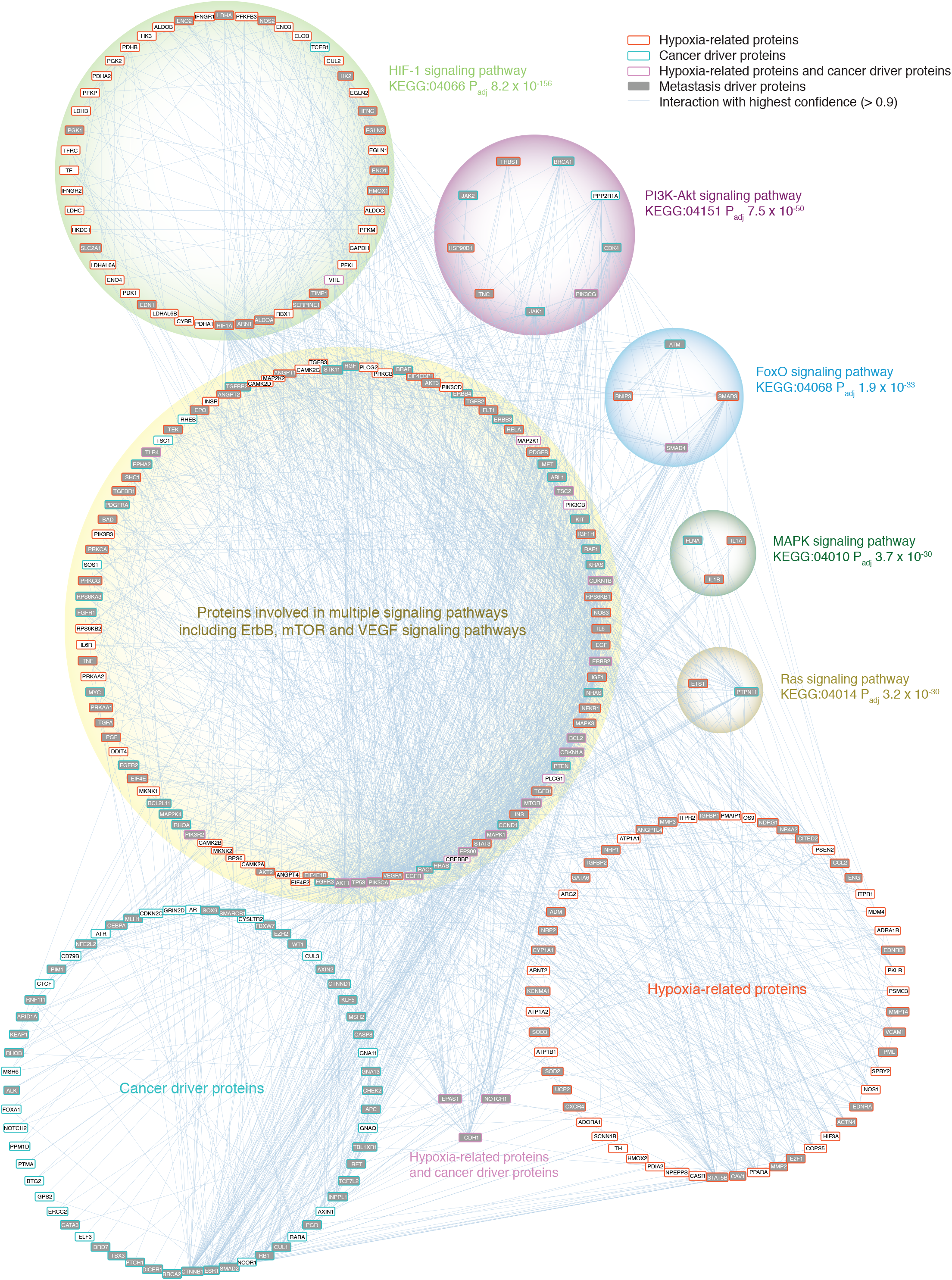
Protein-protein interactome network conformed by hypoxia-related proteins, cancer driver proteins and metastatic driver proteins.

The most significant pathways that make up the PPi network were HIF-1 signaling pathway (P_adj_ = 8.2 × 10^−156^) with 105 nodes, PI3K-Akt signaling pathway (P_adj_ = 7.5 × 10^−50^) with 85 nodes, FoxO signaling pathway (P_adj_ = 1.9 × 10^−33^) with 45 nodes, MAPK signaling pathway (P_adj_ = 3.7 × 10^−30^) with 60 nodes, and Ras signaling pathway (P_adj_ = 3.2 × 10^−30^) with 54 nodes (Table S25). Additionally, top ten proteins with the highest degree centrality in the HIF-1 signaling pathway were AKT1, PIK3CA, HIF1A, VEGFA, EGFR, EP300, CREBBP, STAT3, MAPK1 and INS; in the PI3K-Akt signaling pathway were TP53, AKT1, PIK3CA, VEGFA, EGFR, RAC1, HRAS, CCND1, MAPK1 and INS; in the FoxO signaling pathway were AKT1, EGFR, HRAS, EP300, CREBBP, STAT3, CCND1, MAPK1, INS and TGFB1; in MAPK signaling pathway were TP53, AKT1, EGFR, RAC1, HRAS, MAPK1, INS, TGFB1, NFKB1 and MAPK3; in the Ras signaling pathway were AKT1, PIK3CA, VEGFA, EGFR, RAC1, HRAS, MAPK1, PTPN11, INS and PLCG1. Lastly, the top ten proteins with the highest degree centrality in the entire PPi network were TP53, AKT1, PIK3CA, HIF1A, VEGFA, EGFR, RAC1, HRAS, EP300 and CREBBP.

On the other hand, the Human Cancer Metastasis Database (HCMDB) is an integrated database designed to analyze large scale expression data of cancer metastasis ^59^. Our PPi network was conformed by 179 (62%) metastatic driver proteins (Figure 7) (Table S26). The top ten-metastasis driver proteins with the highest centrality degree were TP53, AKT1, PIK3CA, HIF1A, VEGFA, EGFR, RAC1, HRAS, EP300 and STAT3.

### Functional enrichment analysis using g:Profiler

g:Profiler searches for a collection of proteins representing pathways, networks, gene ontology (GO) terms and disease phenotypes such as cancer ^60,61^. Figure 8A shows the enrichment map of the 233 hypoxia-related proteins. The most significant GO: molecular functions with Benjamini-Hochberg correction and false discovery rate (FDR) < 0.001 were phosphotransferase activity, kinase activity, enzyme binding, signaling receptor binding and small molecule binding. The most significant GO: biological processes were response to hypoxia, response to decreased oxygen levels, response to oxygen levels, response to abiotic stimulus and response to stress. Lastly, the most significant signaling pathways from the Kyoto Encyclopedia of Genes and Genomes (KEGG) were HIF-1 signaling pathway, ErbB signaling pathway, PI3K-Akt signaling pathway, FoxO signaling pathway and mTOR signaling pathway (Table S27).

**Figure 8.**
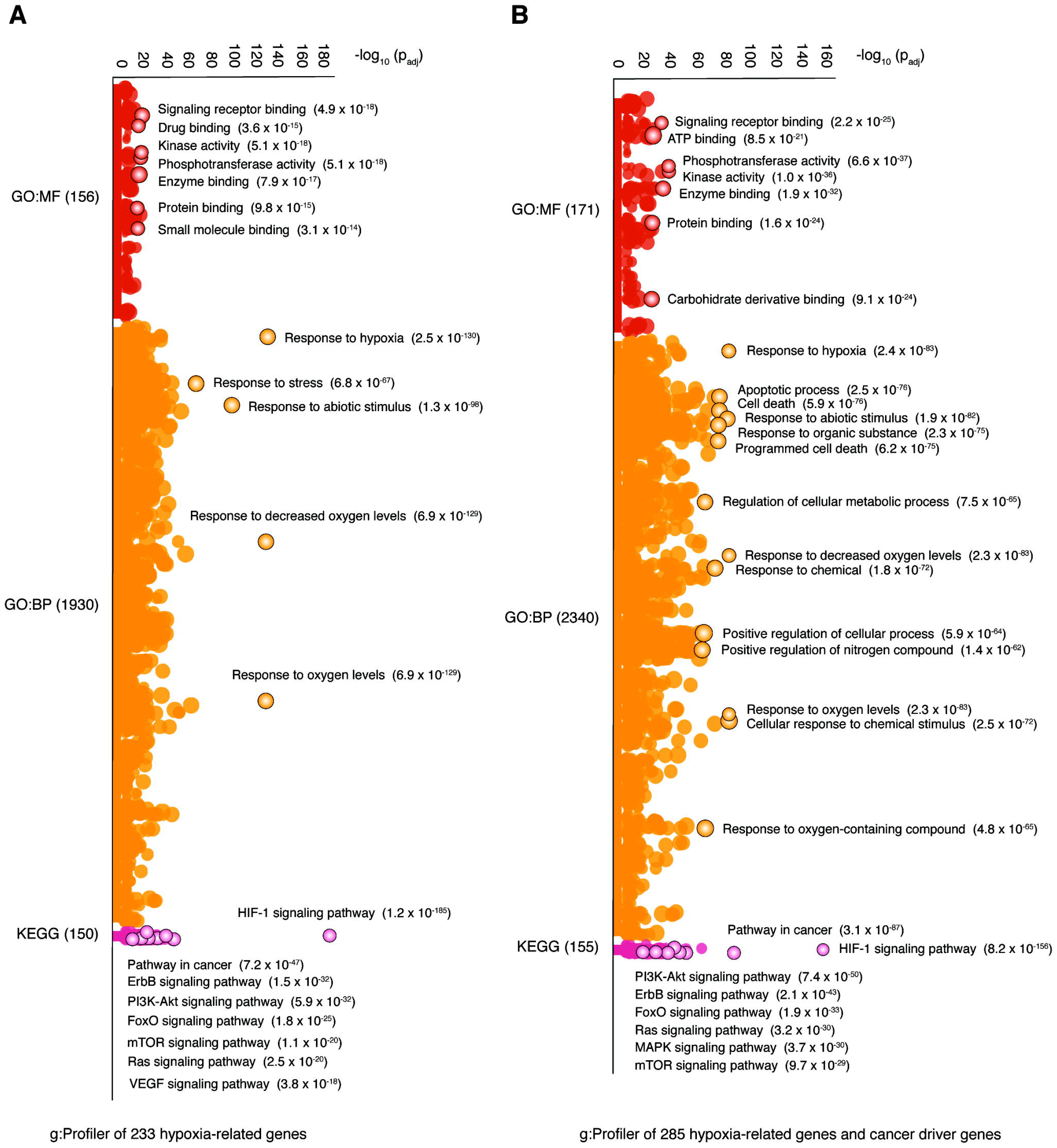
Functional enrichment analysis using g:Profiler. A) Most significant GO molecular functions, GO biological processes and KEGG signaling pathways of the 233 hypoxia-related genes. B) Most significant GO molecular functions, GO biological processes and KEGG signaling pathways of the 285 hypoxic genes and cancer driver genes.

On the other hand, figure 8B shows the enrichment map of 285 proteins conformed by the interaction between hypoxia-related proteins and cancer driver proteins. Phosphotransferase activity, kinase activity, enzyme binding, signaling receptor binding and small molecule binding were the most significant GO: molecular functions. Cellular response to chemical stimulus, response to oxygen levels, response to decreased oxygen levels, response to hypoxia and response to abiotic stimulus were the most significant GO: biological processes. Finally, HIF-1 signaling pathway, pathways in cancer, PI3K-Akt signaling pathway, ErbB signaling pathway and FoxO signaling pathway were the most significant KEGG pathways (Table S28).

### Shortest paths from hypoxia-related proteins to metastasis

We analyzed the 233 hypoxia-related proteins by using CancerGeneNet software in order to find the shortest paths to metastasis according to Iannuccelli *et al* ^62^. We found that 99 (42%) proteins had paths to metastasis, of which, 49 proteins had positive regulation with an average distance of 3.0 and an average path length of 4.0, 16 proteins had negative regulation with an average distance of 3.5 and an average path length of 4.6, and 34 proteins with unknown regulation status had an average distance of 3.2 and an average path length of 4.3. The top ten hypoxia-related proteins with the shortest distance to metastasis were BACH1 (0.8), AKT2 (1.6), AKT1 (1.6), CAMK2B (1.7), EGFR (1.7), MAPK1 (1.7), MAPK3 (1.7), PRKCA (1.7), PRKCB (1.7) and EGF (1.8). Lastly, the shortest paths to metastasis of the 99 hypoxia-related proteins are fully detailed in figure 9 and table S29.

**Figure 9.**
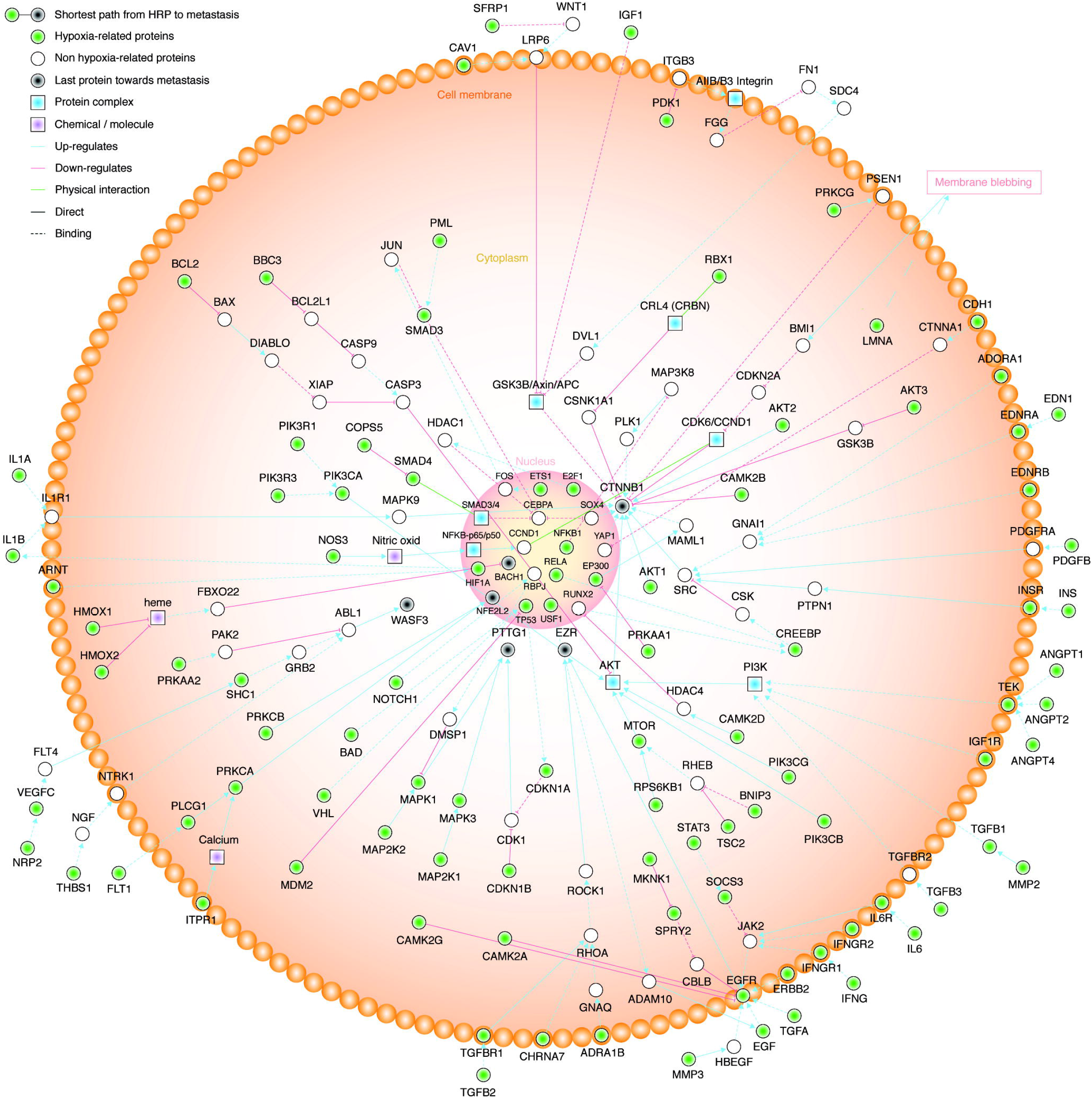
Metastasis signaling. Shortest paths from 99 hypoxia-related proteins to metastasis.

### Hypoxic proteins related to metastasis

Figure 10 shows a Venn diagram of 62 (27%) hypoxic proteins with the highest affinity to metastasis phenotype (Table S30). Additionally, 27 (12%) of them presented a frequency mean of genomic alterations more than the average (> 0.068) across 17 TCGA Pan-Cancer types (Table S1).

**Figure 10.**
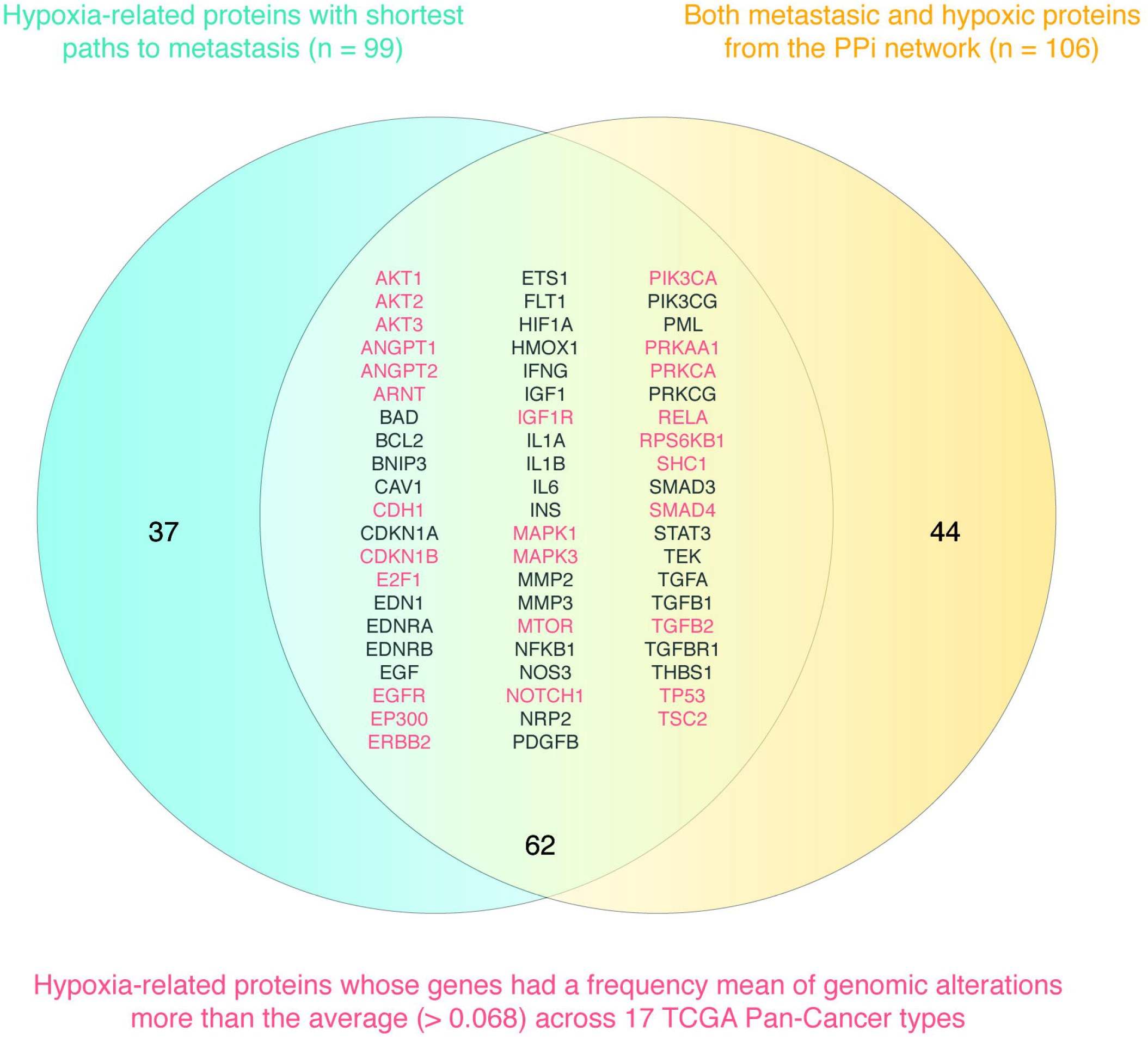
Venn diagram conformed by the hypoxic proteins with shortest paths to metastasis, and the metastatic proteins from the PPi network.

### Clinical trials

Figure 11 shows the current status of clinical trials of drugs focused on metastasis according to the Open Targets Platform ^63^. There were 31 drugs that were being analyzed in 161 clinical trials in 13 of 62 genes. Cancer types with clinical trials focused on metastasis were breast cancer, kidney cancer, liver cancer, colorectal cancer, melanoma and prostate cancer (Figure 11A). *EGFR* presented the highest number of clinical trials in process, recruiting or completed, followed by *FLT1*, *TEK*, *ERBB2*, *IGF1R*, *PIK3CA*, *PIK3CG*, *AKT1*, *AKT2*, *AKT3*, *BCL2*, *NOTCH1* and *TGFBR1* (Figure 11B). Most clinical trials were in phase 2, followed by phase 1 and 4 (Figure 11C). Antibodies and small molecules were the type of drugs focused on metastasis with 86 and 75 clinical trials, respectively (Figure 11D). Inhibition was the most frequent type of action (165), followed by antagonism (5) (Figure 11E). Tyrosine protein kinase EGFR family was the target class with highest number of clinical trials (94), followed by tyrosine protein kinase VEGF family (36), tyrosine protein kinase Tie family (15), enzyme (6), tyrosine protein kinase InsR family (4), AGC protein kinase AKT family (3), Bcl-2 family (1) and TKL protein kinase STKR type 1 subfamily (1) (Figure 11F). Figure 11G diagrams a Sankey plot comparing the number of clinical trials in metastatic drugs analyzed in different cancer types (Table S31 shows the list of clinical trials of drugs focused on metastasis). Lastly, figure S1 and table S32 shows the list of clinical trials in the entire hypoxia-related genes.

**Figure 11.**
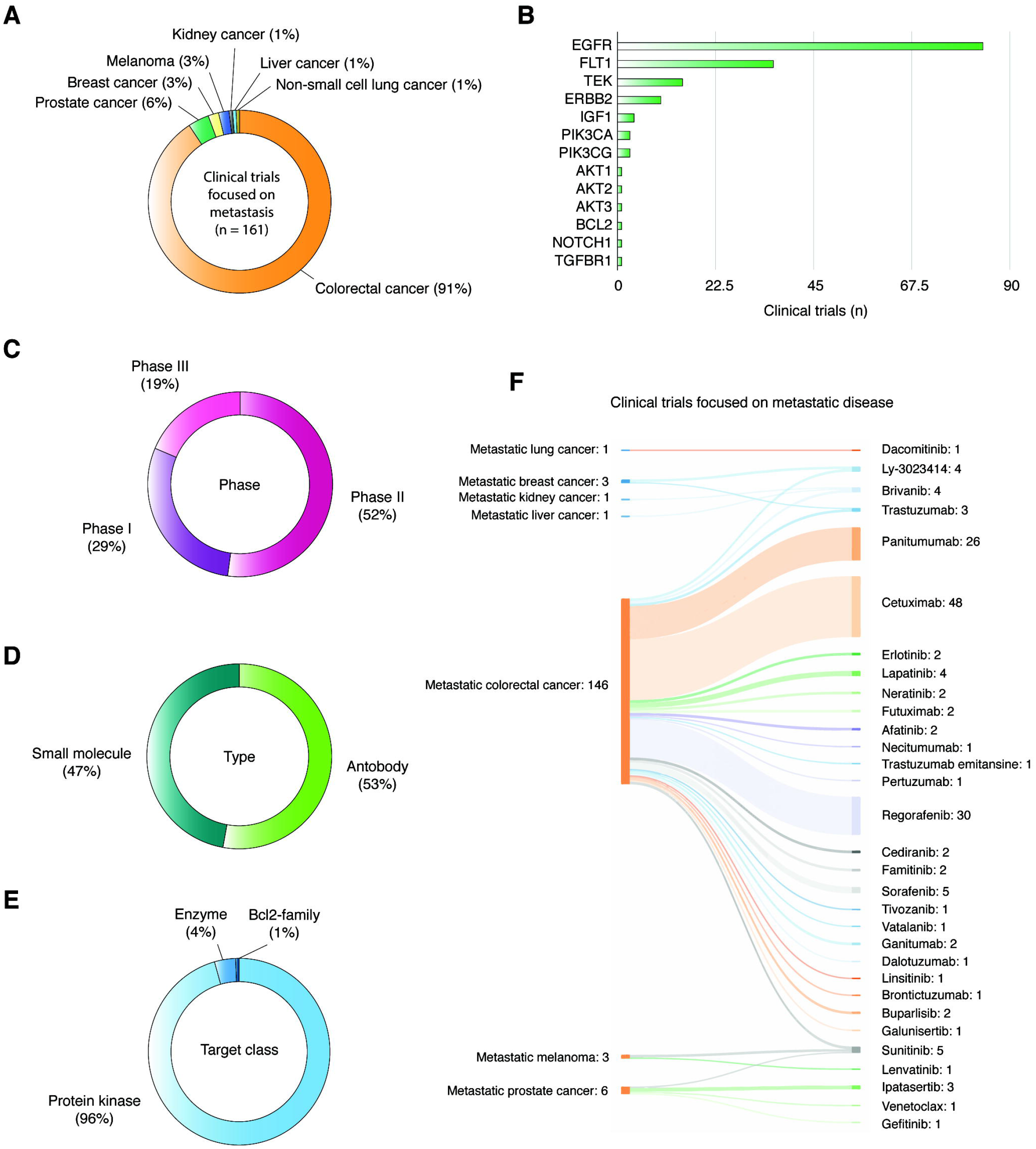
Clinical trials of drugs focused on metastasis. A) Percentage of clinical trials per cancer type. B) Hypoxic genes with highest number of clinical trials on metastasis. C) Phase of clinical trials. D) Type of drugs. E) Action type of drugs. F) Target class. G) Correlation of clinical trials, cancer types and drugs using a Sankey plot.

## DISCUSSION

Hypoxia induces a series of biological changes that contribute to tumorigenesis and metastasis ^24,64^, and are associated with resistance to chemotherapy, radiation therapy, drug therapy and immunotherapy. Therefore, understanding the effect of hypoxia on multi-omics signatures is crucial to improving the outcomes of cancer therapy ^65^.

We analyzed the OncoPrint of 100,643 genomic alterations (233 HRG) of 6,343 individuals across 17 TCGA Pan-Cancer types. Cancer types with the highest frequency mean of genomic alterations were ESCA, BRCA, LUSC, BLCA and LUAD. The genomic alteration with the highest frequency mean in all cancer types was mRNA high because hypoxic tumors trigger an over expression of intracellular signals to adapt to the environment ^12,66,67^. The hypoxia-related genes with the highest frequency mean of mRNA high alteration were *COPS5*, *ELOC*, *ARNT*, *PIK3CA*, *TFRC*, *PSEN2*, *PLCG1*, *USF1*, *E2F1* and *NDRG1*. Lastly, cancer types with the highest frequency mean of mRNA high alteration were ESCA, LUSC, MESO, CESC and BRCA.

Regarding genomic alterations per tumor stage across 17 TCGA Pan-Cancer types, T4 stage showed the highest frequency mean (0.072), followed by T2, T3 and T1. There was a significant difference (Mann-Whitney *U* test, P < 0.001) between T1 and T4 stages. In addition, T4 stage presented the highest frequency means of mRNA high alteration (0.050) and putative driver mutations (0.004). On the other hand, the frequency mean of genomic alterations in HRG was higher in patients with metastasis (0.072) than in patients without metastasis (0.069), with a significant difference (Mann-Whitney *U* test, P < 0.001). That is, the amount of genomic alterations in the HRG increases as the tumor stage and metastasis status evolves.

In order to validate the significant differences of genomic alterations of HRG found in T stages and metastasis status, we analyzed the Buffa hypoxia score across 13 TCGA Pan-Cancer types. The Buffa signature is an approach for deriving signatures that combine knowledge of gene function and analysis of *in vivo* co-expression patterns. Hypoxia scores were estimated by obtaining the mean expression (log2) of 52 hypoxia genes reported by Buffa *et al* ^52,68^. Cancer types with the highest HS mean were HNSC, LUSC, CESC, CRC and BLCA, and the overall HS mean was 0.85. Regarding T stages, T4 presented the highest HS mean (14.5), followed by T2, T3 and T1. There was a significant difference (Mann-Whitney *U* test, P < 0.001) between T1 and T4 stages. On the other hand, the Buffa HS mean in patients with metastasis (8.3) was higher than in patients without metastasis (2.2), with a significant difference (Mann-Whitney *U* test, P < 0.001). In regard to ethnicity, Guerrero *et al*., indicate a need to diversify oncological studies to other populations along with novel strategies to enhance race/ethnicity data recording and reporting ^69^. In this context, Black / African Americans showed highest HS mean (4.4) than Whites and Asians, and Black /African Americans with metastasis showed highest HS mean (12.3) than Whites (8.1) and Asians (−12.3). Notably, the Buffa HS increases as the tumor stage and metastasis status evolves.

In addition to analyzing the genomics and transcriptomics of HRG, we evaluated the human tissue proteome based on tissue microarray-based immunohistochemistry of 175 hypoxia-related proteins across 15 TCGA Pan-Cancer types ^53–55,67^. The AKT3, CAMK2A, PDHA1, PDHB, POLR2A, SMAD4 and SOD3 proteins acted as tumor suppressors, meanwhile, the ADAM8, BBC3, CCL2, CDH1, CLDN3, EDN1, EIF4EBP1, RELA and TN proteins acted as oncogenes according to the expression levels between normal and tumor tissues.

Regarding the hallmarks of cancer proposed by Hanahan and Weinberg ^56^, 23 of 230 HRG showed some of these cancer features. The *TP53*, *AKT1*, *EGFR*, *EPAS1*, *HIF1A*, *MTOR*, *NOTCH1*, *VHL*, *CREBBP* and *ERBB2* genes presented the highest number of associations with these hallmarks. Invasion and metastasis was promoted by *TP53*, *AKT1*, *EGFR*, *EPAS1*, *HIF1A*, *MTOR*, *VHL*, *CREBBP*, *ERBB2*, *NDRG1*, *PIK3CA*, *PLCG1*, *SMAD3*, *ARNT*, *MAPK1*, *PIK3R1*, *ATP1A1*, *CDH1* and *CXCR4*; suppressed by *TP53*, *VHL*, *NDRG1*, *ARNT* and *PIK3R1*; finally, promoted and suppressed by *AKT1* and *CDH1*.

It is important to mention that proteins do not work alone, that is why we performed a protein-protein interactome network analysis to better understand the behavior of hypoxia-related proteins and its correlation with cancer driver proteins and metastasis. The PPi network had 285 nodes conformed by 163 (57%) hypoxic proteins, 98 (34%) cancer driver proteins, and 24 cancer driver proteins with hypoxic signature. Regarding metastasis, our PPi network was conformed by 179 of 285 (62%) metastatic driver proteins. The top ten-metastasis driver proteins with the highest centrality degree were TP53, AKT1, PIK3CA, HIF1A, VEGFA, EGFR, RAC1, HRAS, EP300 and STAT3.

Consequently, we performed two functional enrichment analyses by using g:Profiler ^60^. The first analysis was conformed by the 233 hypoxia-related proteins where the most significant (FDR < 0.001) GO: biological processes were response to hypoxia, response to decreased oxygen levels, response to oxygen levels, response to abiotic stimulus and response to stress. The most significant KEGG signaling pathways were HIF-1, ErbB, PI3K-Akt, FoxO and mTOR. As expected, all terms were related to hypoxia conditions. The second analysis showed the enrichment map of 285 proteins conformed by the interaction between hypoxia-related proteins and cancer driver proteins. The most significant GO: biological processes were cellular response to chemical stimulus, response to oxygen levels, response to decreased oxygen levels, response to hypoxia and response to abiotic stimulus. Finally, HIF-1, PI3K-Akt, ErbB and FoxO were the most significant KEGG signaling pathways.

Another way to analyze the correlation between hypoxia and metastasis was unrevealing the shortest paths from hypoxia-related proteins to metastasis signaling ^62^. We found that 99 of 233 (42%) proteins had paths to metastasis where 49 had positive regulation with an average distance of 3.0, and 16 had negative regulation with an average distance of 3.5. Hypoxic proteins with the shortest distance to metastasis were BACH1, AKT2, AKT1, CAMK2B, EGFR, MAPK1, MAPK3, PRKCA, PRKCB and EGF. Lastly, these 99 hypoxia-related proteins send metastatic signaling through CTNNB1, BACH1, EZR, PTTG, WASF3 and NFE2L2.

After the PPi network analysis and the analysis of the shortest paths to metastasis, we proposed 62 proteins with the highest affinity to metastasis phenotype (Figure 10). Of them, 27 proteins presented a frequency mean of genomic alterations more than the average (> 0.068) across 17 TCGA Pan-Cancer types. Consequently, we observed that 13 of these 62 proteins have 31 drugs studied in 161 clinical trials focused on metastasis. It is important to mention that the percentage of druggable proteins under clinical trials is low and machine-learning analysis could be required in order to improve the selection of drugs against metastasis ^70,71^.

## CONCLUSIONS

Hypoxia and HIF-dependent signaling play an important role in metastasis tumor progression. The hypoxic tumor microenvironment influences both the early and late stages of metastasis. Our study provides a comprehensive view of multi-omics analyses in hypoxia-related genes. Our findings suggest that individuals with metastasis present highest frequency of genomic alterations and hypoxia score across 17 TCGA Pan-Cancer types. The most altered pathways where hypoxic genes are involved are HIF-1, ErbB, PI3K-Akt, FoxO, mTOR and Ras signaling pathways. The analyses of both the PPi network and the shortest paths to metastasis suggest 62 proteins strongly related to metastasis. Finally, there are few drugs anti-metastasis that are being studied in clinical trials so far. Hence, it is imperative to focus our strengths in drug developing and drug discovery to improve precision medicine and find effective therapies anti-metastasis.

## METHODS

### Dataset of hypoxia-related genes

In this study we analyzed a dataset of 233 HRG curated manually. 109 of them conformed the KEGG HIF-1 signaling pathway ^72^, 102 of them were involved in some of the GO terms related to hypoxia: response to hypoxia, cellular response to hypoxia and cellular response to decreased oxygen levels by using g:Profiler ^60^, 52 of them were hypoxia genes according to Buffa *et al* ^52^, and 75 of them underlie high-altitude adaptive phenotypes in populations from the Andean Altiplano, Semien Plateau and the Tibetan Plateau, according to Bigham ^73^.

### Genomic alterations

Genomic alterations (mRNA high, mRNA low, CNV deep deletion, CNV amplification, fusion gene, inframe mutation, truncating mutation and missense mutation) were analyzed in 6,343 tumors (all with tumor stage) from 17 TCGA Pan-Cancer types (BLCA, BRCA, CESC, CRC, ESCA, HNSC, KIRC, LIHC, LUAD, LUSC, MESO, PAAD, PRAD, SKCM, STAD, TGCT and THCA). Genomics and clinical data related to tumor stage (T1-T4) and metastasis status (M0-M1) was taken from the Genomics Data Commons of the National Cancer Institute (https://portal.gdc.cancer.gov/) and the cBioPortal (http://www.cbioportal.org/) ^50,51^.

### Hypoxia score

The Buffa hypoxia score was analyzed in 5,249 tumors from 13 TCGA Pan-Cancer types (HNSC, LUSC, CESC, CRC, BLCA, SKCM, KIRC, LUAD, PAAD, BRCA, LIHC, PRAD and THCA) ^52^. An approach for deriving signatures that combine knowledge of gene function and analysis of *in vivo* co-expression patterns was used to define a common hypoxia signature in cancer. Hypoxia scores were estimated by obtaining the mean expression (log2) of 52 hypoxia genes reported by Buffa *et al* ^52,68^.

Data related to Buffa hypoxia score, tumor stage, metastasis status and ethnicity (black / African American, white and Asian) was taken from the Genomics Data Commons and the cBioPortal ^50,51^. Lastly, the Mann-Whitney *U* test was performed to determine significant differences between hypoxia score and clinical data.

### Protein expression analysis

The Human Protein Atlas (https://www.proteinatlas.org/) explains the diverse molecular signatures of proteomes in the human tissue based on tissue microarray-based immunohistochemistry ^53,54,74,75^. We compared the protein levels (high, medium, low, non-detected) of our 175 proteins between normal tissues and tumor tissues from 15 different cancer types. Lastly, we obtained a list of proteins with potential interaction as oncogenes and tumor suppressor genes.

### Protein-protein interactome network

The PPi network with a highest confidence cutoff of 0.9 and zero node addition was created using the String Database, which takes into account predicted and known interactions ^57,76,77^. The degree centrality of a node means the number of edges the node has to other nodes in the network. Metastasis driver proteins that interact with the hypoxia-related proteins were taken from the Human Cancer Metastasis Database (https://hcmdb.i-sanger.com/). HCMDB is an integrated database designed to analyze large scale expression data of cancer metastasis ^59^. The network visualization was analyzed through the Cytoscape software ^58^. Hypoxia-related proteins, cancer driver proteins, metastasis driver proteins, and significant signaling pathways (P < 0.001) were differentiated by colors in the PPi network.

### Functional enrichment analysis using g:Profiler

The enrichment analysis gives scientists curated interpretation of gene/protein lists from genome-scale experiments ^60^. The hypoxia-related proteins and cancer driver proteins were analyzed by using g:Profiler (https://biit.cs.ut.ee/gprofiler/gost) in order to obtain significant annotations (Benjamini-Hochberg (FDR) < 0.001) related to GO terms (molecular functions and biological processes), and KEGG signaling pathways ^60,72^.

### Shortest paths from hypoxia-related proteins to metastasis

CancerGeneNet (https://signor.uniroma2.it/CancerGeneNet/) is a resource that links genes that are frequently mutated in all cancer types to cancer phenotypes. This resource is based on the annotation of experimental information that permits to embed the cancer genes into the cell network of causal protein relationships curated in SIGNOR ^78^. Consequently, this bioinformatics tool allows to infer likely paths of causal interactions linking cancer associated genes to cancer phenotypes such as metastasis ^62^. Iannuccelli *et al* explained that the shortest paths between a specific gene and cancer phenotypes was programmatically implemented using the shortest path function of the R *igraph*, obtaining a distance score and a path length score ^62,79^. Hence, we analyzed the shortest paths from our hypoxia-related proteins to metastasis to better understand the association to this hallmark of cancer ^62,80^.

### Clinical trials

The Open Targets Platform (https://www.targetvalidation.org) is comprehensive and robust data integration for access to and visualization of potential drug targets associated with several cancer types and metastasis. Additionally, this platform shows all drugs in clinical trials associated with biomarkers or cancer genes, detailing its phase, type of drug, action type, and target class ^63^.

## Acknowledgments

Universidad UTE (Ecuador) and the Latin American Society of Pharmacogenomics and Personalized Medicine (SOLFAGEM) supported this research.

## Author contribution

ALC and PGR conceived the subject and the conceptualization of the study. ALC wrote the manuscript. CPyM supervised the project. ALC and CPyM did founding acquisition. PGR, SG, JMGC, AKZ, IAC, APV, VY, DCB and PEL did data curation and supplementary data. SG, EOP and NV gave conceptual advice and valuable scientific input. Finally, all authors reviewed and approved the manuscript.

## Data availability statement

All data generated or analyzed during this study are included in this published article (and its Supplementary Information files).

